# Identification of bacteria by poly-aromatic hydrocarbons biosensors

**DOI:** 10.1101/2021.11.27.470193

**Authors:** Yaniv Shlosberg, Yair Farber, Salah Hasson, Valery Bulatov, Israel Schechter

**Affiliations:** Schulich Faculty of Chemistry, Technion, Haifa 3200003, Israel; Quality and Reliability Engineering Department, Kinneret Academic College, Zemach 1513200, Israel; Grand Water Research Institute, Technion, Haifa 3200003, Israel

**Keywords:** Identification, Bacteria, Polyaromatic hydrocarbons, Perylene, Synchronous fluorescence

## Abstract

Human health is consistently threatened by different species of pathogenic bacteria. To fight the spread of diseases, it is important to develop rapid methods for bacterial identification. Over the years, different kinds of biosensors were developed for this cause. Another environmental risk are poly-aromatic hydrocarbons (PAHs) that may be emitted from industrial facilities and pollute environmental water and soil. One of the methods for their purification is conducted by the addition of bacteria that can degrade the PAHs, while the bacteria itself can be filtrated at the end of the process. Although many studies reported monitoring of the PAHs degradation by fluorescence, not much attention was dedicated to studying the influence of the PAHs on the intrinsic fluorescence of the degrading bacteria. In this work, we apply synchronous fluorescence (SF) measurements to study the ability of the 5 PAHs: 9-Antracene carboxylic acid (9ACA), Pyrene, Perylene, Pentacene, and Chrysene to interact with bacteria and change its fluorescence spectra. We show that upon incubation of each PAH with the bacterium *E*.*coli* only the 2 PAHs 9ACA and Perylene cause an intensity decrease in the emission at λ = 300 – 375 nm, which derives from the emission of Tyrosine and Tryptophane (TT). Also, we show that upon incubation of 9ACA and Perylene with 5 different pathogenic bacteria, the intensity increase or decrease in the TT emission is unique to each bacterial species. Based on this observation, we suggest that the PAHs 9ACA and Perylene can be utilized as biosensors for bacterial identification.

## Introduction

A major risk that has always threatened the survival of mankind is the pathogenic bacteria. To fight this threat and prevent the spread of diseases, many efforts were done over the years to develop analytical methods for bacterial identification. A traditional method that is used to distinguish between gram-negative and positive bacteria can be done by gram staining followed by microscopic observation. [1] This method can be very useful for initial diagnostic because it is simple and rapid. However, it cannot identify the species of the tested bacterium. Specific identification can be done conducted by various advanced microscopic methods, such as fluorescence microscopy. [2] To enable the specificity of the detection under the microscope, immune fluorescence approaches can be applied, in which antigens in the external layer of the bacterium are targeted by antibodies that are conjugated with a fluorescent molecule. [3] Another possibility that is in many cases less expensive is to use a non-fluorescent antibody to target the analyte and a secondary fluorescent antibody that bind to the primary one. [4] An efficient method that can count the number of cells and detect the presence of antibodies is Flow cytometry. [5] In this method, the cells are inserted into a microfluidic device that enables to detect their fluorescence one by one by one using a laser as a strong light source. A more advanced method is flow cytometry imaging that can simultaneously integrate both flow cytometry and fluorescence microscopic observation in multi-channels. [6] Another approach for bacterial identification is electron microscopy. [7] Electron microscopy setups enable to get a high imaging resolution of the bacterium which is in many cases sufficient for its identification based on its shape, size, and other visual properties. Although optical and electron microscopy can successfully identify bacteria, they are mostly expensive and non-portable. Therefore, they may not be optimal for wide usage or outdoor environmental monitoring of water sources. A different approach for bacterial monitoring is the utilization of electrochemical methods. [8, 9] Bacteria are composed of both components that are capable and incapable of charge transfer. This dual nature enables the detection of bacteria by Faradaic electrochemical methods such as cyclic voltammetry that is based on the oxidation/reduction of constitutes at the external layer of the bacterium and secreted metabolites. Also, Non-Faradaic methods such as electrochemical impedance spectroscopy can be used to detect bacteria based on their resistance. [10, 11] In many cases, the differences between the electrochemical signal of bacterial species are not sufficient to differentiate between them. Therefore, a specific targeting method should be conducted prior to electrochemical detection. This can be done by depositing conductive polymers for bacterial imprinting on electrodes. [12] In this approach, conductive monomers are polymerized together with the bacterial analyte to form a thin film with trapped bacteria at its surface. Different treatments such as heating can destruct the bacterial structure to release them from the surface without harming the polymer itself. [12] The release remains cavities in the surface of the polymer with the shape and size of the bacterium that was once trapped there. The film with the cavities can be used as a specific biosensor for bacterial targeting, while its deposition on the electrode enables to make a fast electrochemical detection. [8, 9] An additional efficient method for targeting bacteria for electrochemical detection is the utilization of Aptamers. [13] These short DNA segments or peptides can be designed to selectively bind components in the bacteria. Attachment of the Aptamers to the electrode surface enables their electrochemical identification. One of the advantages of using the targeting methods of Aptamers binding and imprinting bacterial imprinting-based technologies is the possibility to use the same sample for both electrochemical and optical measurements. [14] over the years, many optical spectroscopic methods were developed for bacterial identification. Among the leading methods in the field is Fourier transform infrared spectroscopy (FTIR) and surface-enhanced Raman Spectroscopy (SERS). [15–19] One of the great advantages of these technologies derives from their ability to identify bacteria without using any labeling components. The different bacterial species are composed of building blocks such as lipids, proteins, sugars, etc. However, the different quantities of these biomaterials can be a base for bacterial identification. In some cases, the spectra of different species are very similar, while no significant differences can be observed. One of the ways to overcome this challenge is by making a big number of repetitions and applying chemometric methods. [20] An additional spectroscopic method that relies on different quantities of bacterial ingredients is near-infrared spectroscopy. [21–25] A major advantage of this method derives from its ability to penetrate glass, which allows to easily measure bacteria in their cultivation vessel without damaging its sterilization. An additional non-invasive method is Terahertz spectroscopy. [26] The in the Terahertz wavelengths range, the absorption coefficients of the biomaterials in bacteria are low. This enables the irradiation to penetrate and collect spectral data from deeper areas of the bacterial cell without harming it. [26] A different method that is efficient but destructive for the cells is laser-induced breakdown spectroscopy (LIBS). [27–33] This technique can differentiate between species based on the differences in the quantities of their atoms.

Another leading approach for bacterial identification is the utilization of fluorescence spectroscopy. [34–46] Microorganisms are consist of several components such as the amino acids Tyrosine, Tryptophane, and Phenylalanine, or metabolites such as NADPH that can be detected based on their intrinsic fluorescence. [47, 48] DNA also fluoresce under UV-VIS irradiation, however, its native emission is usually too low to be detected. Amplification of DNA fluorescence can be done by labeling it with fluorescent dyes such as SYBR that amplify its emission intensity. [49] While performing fluorescence measurements, choosing the scanning method and parameters may be highly important. 2D - fluorescence maps can collect the full spectral data of the analyte, however, the long exposure to the UV-Vis irradiation may inflict photodamage if it lasts too long. [50, 51] This problem can be overcome by using a CCD detector instead of a photomultiplier, however, such detectors may lower the sensitivity of the detection. [52] The most frequently used scanning method is the emission scan in which λ_Ex_ is fixed and λ_Em_ is variable. Using this scan mode for most bacteria will produce broad peaks that are hard to analyze and be used for bacterial identification unless assisted by chemometrics methods [35, 39–46]. An improved scanning mode for bacterial measurements that narrows the widths of the peaks is synchronous fluorescence (SF) [34] in which both λ_Ex_ and λ_Em_ are variable with a constant Δλ between them [53–55]. Several SF studies were conducted, reporting the identification of *Pseudomonas* colonies directly from its cultivation medium by optic fiber. [38] SF was also used to diagnose complex matrixes such as infection in human urine [36] and spoilage of chicken breast fillets. [37]

One of the methods for environmental purification of water and soil from the pollution of Poly-aromatic hydrocarbons (PAHs) is the utilization of bacterial species that can degrade the PAHs. [56–58] This process can be efficiently monitored based on fluorescence measurements of the PAHs. [59] Although many studies were conducted about this purification approach, not much is known about the biological reaction that occurs in the bacterial cells as a consequence of interaction with different PAHs. In this work, we test the ability of 5 PAHs that are consisted of 2 – 5 aromatic rings to apply as biosensors for the identification of 5 bacterial pathogens. We show that the PAHs Pyrene, Pentacene, and Chrysene do not cause any change to the fluorescence spectra of the bacterium EC, while 9ACA and Perylene decrease their fluorescence intensity at the emission wavelength of tryptophan and tyrosine (TT). Upon incubation of 9ACA and Perylene with 5 different pathogenic bacteria, a unique intensity decrease, or increase is obtained for each species, that can be used as a fingerprint for its identification.

## Materials and Methods

### Bacterial cultivation and sample preparation

6 pathogenic bacterial species *Escherichia coli CN13* (a nalidixic acid-resistant strain) (EC), (ATCC 700609), *Bacillus subtilis* (non-sporulating strain) (BS) (Difco, USA), *Deinococcus radiodurans* (DR) (ATCC 13939), *Enterobacter cancerogenus* (ENC) (isolated from drinking water and identified by Biolog System, USA), *Citrobacter freundii* (CF) (isolated from drinking water and identified by Biolog System, USA) were plated on nutrient agar dishes and incubated at 37 °C for 24 h. After incubation, the bacteria colonies were removed by a spatula and suspended into a small 1 mL saline solution (0.9% NaCl). Prior to fluorescence measurements, normalization of cells quantities was done by dilution of the suspension with saline followed by spectrophotometric measurements (UV-Visible spectrometer HP 8453) to obtain OD_600nm_ = 0.05 according to Janke et al. [60]

### Incubation of PAHs with bacteria

9-Antracene carboxylic acid (9ACA), Pyrene, Perylene, Pentacene, and Chrysene (Merck) were first dissolved in Acetone to form a concentration of 5 mM. 1 mL of each solution was diluted by 9 mL of saline solution (0.9% NaCl). 2 mL of each PAH suspension was incubated for 1 h with 2 mL of a bacterial solution.

### Fluorescence measurements

Fluorescence spectra were measured using a fluorescence spectrophotometer (Aminco Bowman II Luminescence spectrofluorimeter, Thermo, USA). SF measurements were done with an emission resolution of 1 nm. In all measurements, the bandwidths for both excitation and emission were 4 nm.

### Mathematical Analysis and plotting

Chemical structures were drawn using ChemDraw software (Perkin Elmer). All plotting and mathematical analysis were done by OriginPro software.

## Results and Discussion

### Selection of PAHs that can be utilized for sensing EC

In our previous studies, [34] we showed that the amino acids Tyr and Trp (TT) in bacteria can be efficiently quantitated by synchronous fluorescence (SF), while the optimal Δλ value was 60 nm. We wished to explore whether the interaction of bacteria with different PAHs can affect the fluorescence spectra of TT. To estimate the spectral overlap between bacteria and various PAHs, we measured the SF spectra (Δλ = 60 nm, λ_emission_ = 250 – 500 nm) of the bacterium *Escherichia coli CN13* (EC) and the 5 PAHs, 9-Antracene carboxylic acid (9ACA), Pyrene, Perylene, Pentacene, and Chrysene (Scheme. 1), whose molecular structure is consisted of 3 – 5 aromatic rings in different organizations. (Fig.2).

**Fig. 1.**
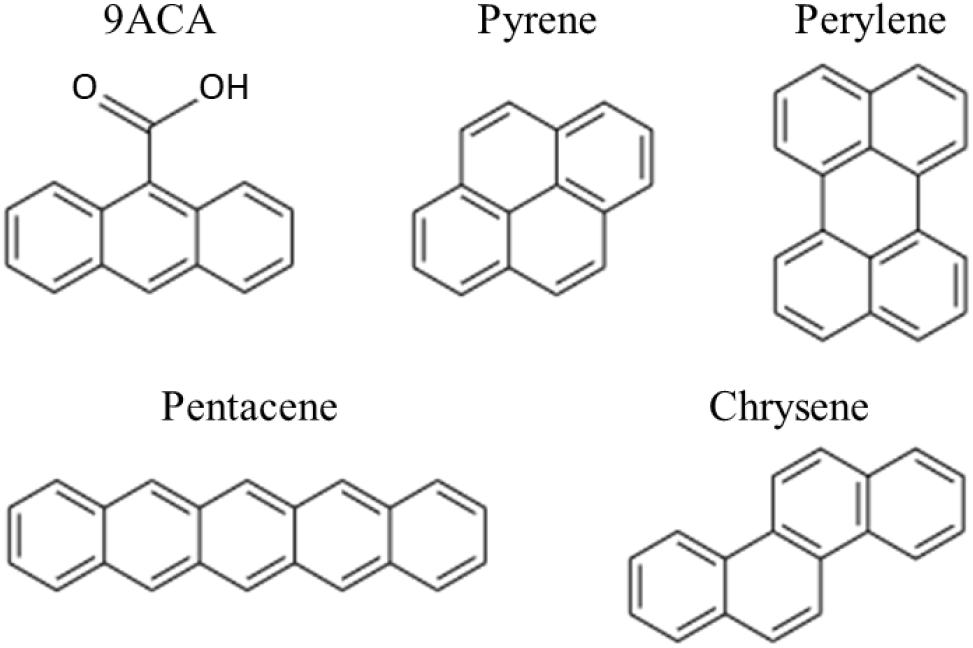
Molecular Structure of the 5 PAHs: 9-Antracene carboxylic acid (9ACA), Pyrene, Chrysene, Perylene, and Pentacene.

**Fig. 2.**
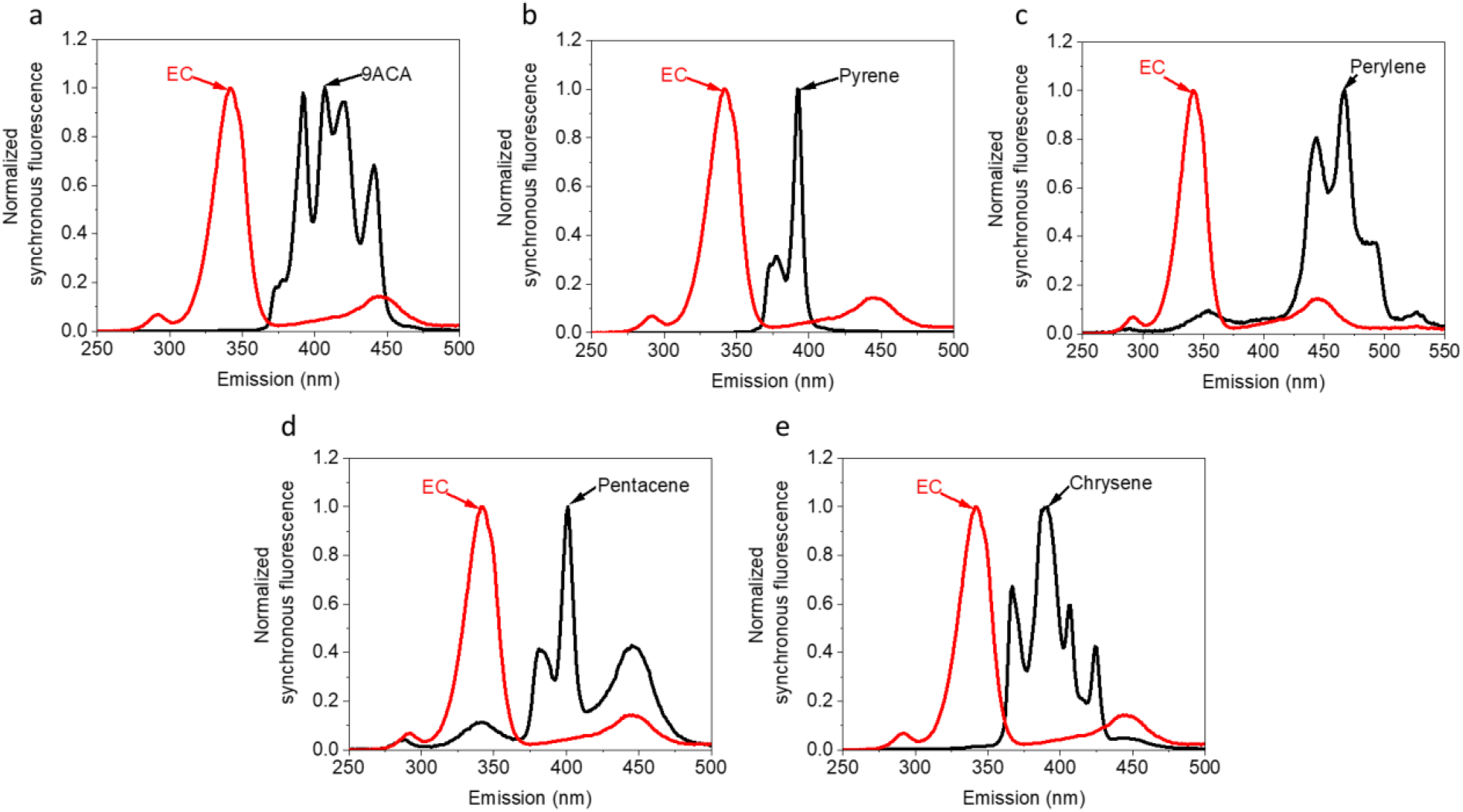
Estimation of the spectral overlap in the SF spectra of EC and 3-5 membered rings PAHs. SF spectra (Δλ = 60 nm) of EC and the 5 PAHs were measured and plotted together to evaluate their spectral overlap. **a** EC + 9ACA. **b** EC + pyrene. **c** EC + Perylene. **d** EC + Pentacene. **e** EC + Chrysene.

The spectral overlap between the normalized spectra of EC and the 3-4 membered rings PAHs: 9ACA, Pyrene, and Chrysene were very small, while these PAHs did not have any significant fluorescence contribution at 342nm (the maxima of the EC spectra). The spectra of the 5 membered ring Pentacene and Perylene have a peak with maxima at 345 and 350 nm respectively. However, these peaks are small and do not process the main fluorescence contribution of the PAHs that is mostly between 400 – 500 nm. The intensity ratio percentage between the normalized spectra of Pentacene, Perylene, and EC at 342 nm was 11 and 7.1 % respectively. The obtained results show that upon incubation of EC with PAHs, the 3 and 4 membered rings 9ACA, Pyrene, and Perylene will probably not change the emission of TT around 300 – 350 nm unless they interact with each other. In the case of the 5 membered rings Pentacene and Chrysene, the spectral overlap must be taken into concern while screening for an interaction that may inflict a spectral change.

To explore whether an interaction between PAHs and EC that changes the SF spectra occur. EC was incubated with each of the 5 PAHs: 9-Antracene carboxylic acid (9ACA), Pyrene, Perylene, Pentacene, and Chrysene,) for 1 h. Following the incubation, the SF of the mixtures were measured (Δλ = 60 nm, λ_emission_ = 250 – 500 nm) and their spectra were compared with the mathematical sum of their unmixed (pure) spectra. No spectral change was observed between the real mixtures and the mathematical sum of the SF spectra of EC and Pyrene, Pentacene, or Chrysene. However, when EC was incubated with 9ACA, and Perylene, the emission intensity of TT (300 – 375 nm) was decreased to 74.7 and 73.1 % respectively (Fig. 3a and 4a). The reason why only 2 of the 5 PAHs can spectrally interact with EC is not fully understood. The structural difference between 9ACA and Perylene to the other 3 PAHs is that they consist of less than 4 conjugated aromatic rings. (9ACA consist of 3 conjugated rings, and Perylene consists of 2 pairs of 2 conjugated rings). We hypnotize that perhaps this structural difference may influence the ability of the PAHs to be internalized by the cells or stick to their outer surface.

**Fig. 3.**
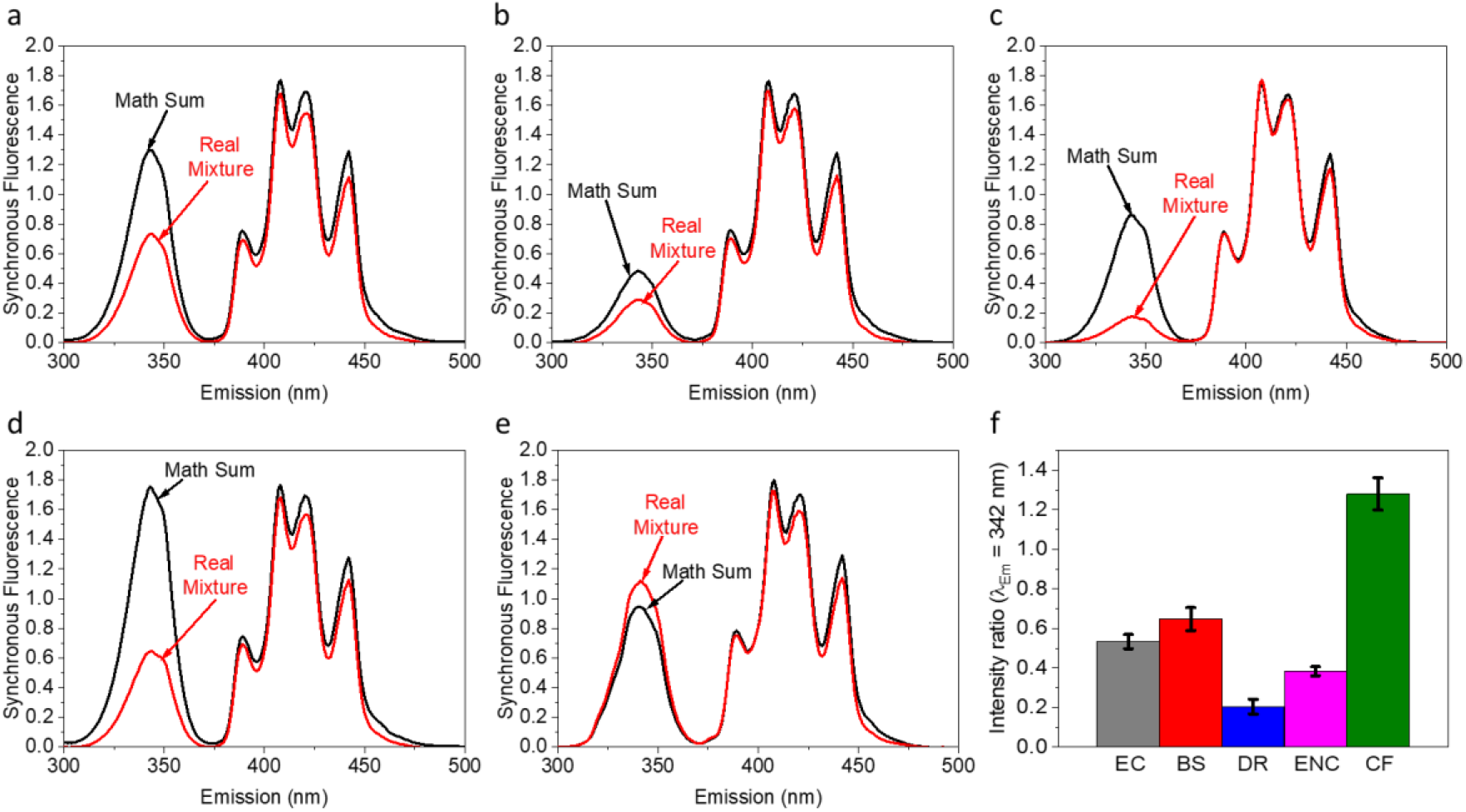
Differentiation of pathogenic bacterial species based on interaction with 9ACA. SF spectra (Δλ = 60 nm) were measured for pure 9ACA + EC, BS, DR, ENC, and CF and the real mixtures of 9ACA + each of the bacterial species after incubation of 1 h. The mathematical sum of 9ACA + each bacterium (black) was plotted vs. the SF spectra of the real mixtures (red). The intensity ratio between the real mixtures and the mathematical sum at the maxima of the peak that derives from the emission of TT in the bacteria (λ=342 nm) was calculated. **a** 9ACA + EC, **b** 9ACA + BS, **c** 9ACA + DR, **d** 9ACA + ENC, **e** 9ACA + CF. **f** The ratio between the real mixtures and the mathematical sum at λ=342 nm. The Error bars represent the standard deviation over 3 independent repetitions.

**Fig. 4.**
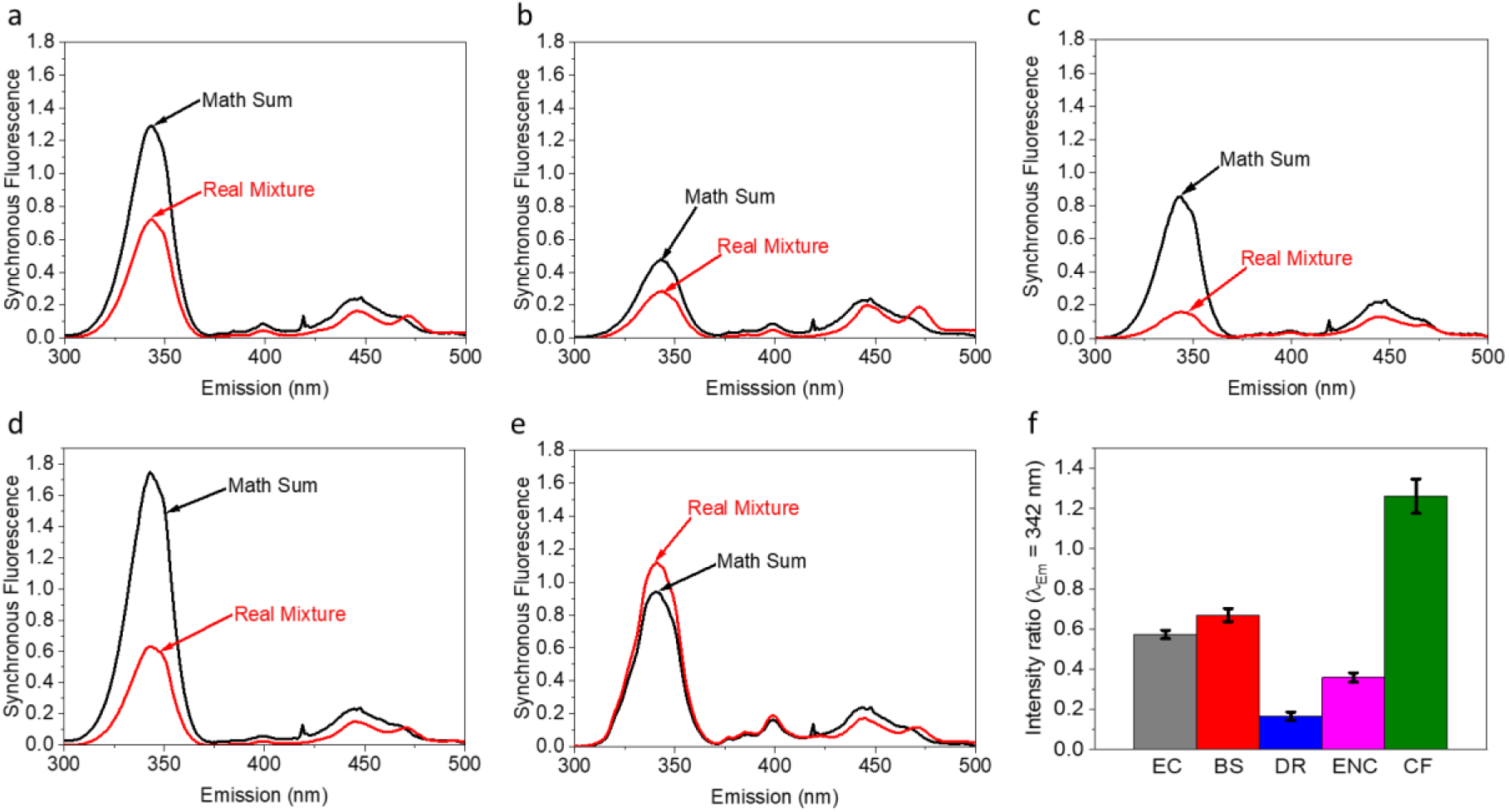
Differentiation of pathogenic bacterial species based on interaction with Perylene. SF spectra (Δλ = 60 nm) were measured for pure Perylene + EC, BS, DR, ENC, and CF and the real mixtures of Perylene + each of the bacterial species after incubation of 1 h. The mathematical sum of Perylene + each bacterium (black) was plotted vs. the SF spectra of the real mixtures (red). The intensity ratio between the real mixtures and the mathematical sum at the maxima of the peak that derives from the emission of TT in the bacteria (λ=342 nm) was calculated. **a** Perylene + EC, **b** Perylene + BS, **c** Perylene + DR, **d** Perylene + ENC, **e** Perylene + CF. **f** The ratio between the real mixtures and the mathematical sum at λ=342 nm. The Error bars represent the standard deviation over 3 independent repetitions.

### Differentiation of pathogenic bacterial species based on interaction with 9ACA

Next, we wished to explore whether 9ACA can also cause a spectral change in other pathogenic bacteria and if this change differs among the species. 9ACA was incubated for 1 h with each of the 5 bacterial species: *Escherichia coli CN13* (EC), *Bacillus subtilis* (BS) *Deinococcus radiodurans* (DR), *Enterobacter cancerogenus* (ENC), and *Citrobacter freundii* (CF). SF spectra of the pure components and their mixtures (Δλ = 60 nm, λ_emission_ = 250 – 500 nm) were measured. (Figs. 3). The obtained results showed that mixing ACA with EC, BS, DR, and ENC has decreased the emission intensity of TT by different coefficients. Interestingly, an opposite trend of an intensity increase was obtained by CF. In all the measurements, the intensity at λ_emission_ = 375 – 500 nm) was slightly decreased. However, the intensity of this decrease was similar in all the measurements and did not show any unique pattern that can be useful for discrimination between the species. The obtained results show it is possible to differentiate between the species based on the ratio between the real mixtures and the mathematical sum of their pure ingredients.

### Differentiation of pathogenic bacterial species based on interaction with Perylene

Following the observation that bacterial species can be differentiated based on the interaction with 9ACA, we next wished to explore whether a similar result can be obtained for Perylene. SF spectra of Perylene, the 5 bacterial species (shown in Fig.3), and their real mixtures were measured. The mathematical sum of Perylene + each bacterium was plotted against the spectra of their real mixtures (Fig. 4). Like the results obtained for 9ACA, mixing perylene with EC, BS, DR, and ENC have inflicted a decrease in the emission of TT, while the interaction with CF has increased it. The intensity change was different for each mixture allowing to discriminate the species based on the intensity ratio between the spectra of the real mixtures and their mathematical sums. In both reactions with 9ACA and Perylene, the bacterial species could be organized based on the ratio (λ_Em_ = 342 nm of the Real mixture / λ_Em_ = 342 of the mathematical sum) by the same order in which DR < ENC < EC < BS < CF. This may imply that both PAHs have triggered a similar reaction in the cells. Another interesting spectral pattern was observed for the peak λ_emission_ = 425 – 500 nm. The mathematical sum of all mixtures showed a peak with maxima at 450 nm with a small shoulder around 470 nm that derives from the emission of both Perylene and NADH in bacteria [47]. In the real mixtures of Perylene with all the bacterial species, an intensity decrease around 450 nm and an increase around 470 nm was observed. These results imply that a direct or indirect reaction that involves both Perylene and NADH might have occurred. As the NADH pools of bacteria are in the cytoplasm, we postulated that perylene can enter bacterial cells. We suggest that the reduction in the TT emission may result from a decrease in the viability of the bacterial cells that is caused by the toxic effects of the PAHs. Another possibility may be that the PAHs intercalate in the bacterial DNA and impair the translation process. In the case of the intensity increase of CF, we suggest that this bacterium can degrade the PAHs using them as a carbon source that induces translational activity. Another possibility can be that CF cells can sense the PAHs and, as a response, promote the biosynthesis of proteins that participate in the degradation of the PAH.

## Acknowledgments

This work was supported by the Israeli Ministry of Science and Technology, by the Office of the Chief Scientist, Israeli Ministry Economy, and by Technion—Israel Institute of Technology (VPR fund).

## Declarations

### Availability of data and material

Not applicable

### Code availability

Not applicable

### Authors’ contributions

YS and IS conceived the idea. YS, SH, VB and IS designed the experiments. YS and YF performed the main experiments. YS wrote the paper. IS provided all funding for the project. IS supervised the entire research project.

